# Genome-wide assessment of *Mycobacterium tuberculosis* conditionally essential metabolic pathways

**DOI:** 10.1101/534289

**Authors:** Yusuke Minato, Daryl M Gohl, Joshua M. Thiede, Jeremy M. Chacón, William R. Harcombe, Fumito Maruyama, Anthony D. Baughn

**Author notes:** Address correspondence to Anthony D. Baughn.

## Abstract

Better understanding of essential cellular functions in pathogenic bacteria is important for the development of more effective antimicrobial agents. We performed a comprehensive identification of essential genes in *Mycobacterium tuberculosis*, the major causative agent of tuberculosis, using a combination of transposon insertion sequencing (Tn-seq) and comparative genomic analysis. To identify conditionally essential genes by Tn-seq, we used media with different nutrient composition. Although many conditional gene essentialities were affected by the presence of relevant nutrient sources, we also found that the essentiality of genes in a subset of metabolic pathways was unaffected by metabolite availability. Comparative genomic analysis revealed that not all essential genes identified by Tn-seq were fully conserved within the *M. tuberculosis* complex including some existing anti-tubercular drug target genes. In addition, we utilized an available *M. tuberculosis* genome-scale metabolic model, iSM810, to predict *M. tuberculosis* gene essentiality *in silico*. Comparing the sets of essential genes experimentally identified by Tn-seq to those predicted *in silico* reveals the capabilities and limitations of gene essentiality predictions highlighting the complexity of *M. tuberculosis* essential metabolic functions. This study provides a promising platform to study essential cellular functions in *M. tuberculosis*.

**Author Summary:** *Mycobacterium tuberculosis* causes 10 million cases of tuberculosis (TB) resulting in over one million deaths each year. TB therapy is challenging because it requires a minimum of six months of treatment with multiple drugs. Protracted treatment times and the emergent spread of drug resistant *M. tuberculosis* necessitate the identification of novel targets for drug discovery to curb this global health threat. Essential functions, defined as those indispensable for growth and/or survival, are potential targets for new antimicrobial drugs. In this study, we aimed to define gene essentialities of *M. tuberculosis* on a genome-wide scale to comprehensively identify potential targets for drug discovery. We utilized a combination of experimental (functional genomics) and *in silico* approaches (comparative genomics and flux balance analysis). Our functional genomics approach identified sets of genes whose essentiality was affected by nutrient availability. Comparative genomics revealed that not all essential genes were fully conserved within the *M. tuberculosis* complex. Comparing sets of essential genes identified by functional genomics to those predicted by flux balance analysis highlighted gaps in current knowledge regarding *M. tuberculosis* metabolic capabilities. Thus, our study identifies numerous potential anti-tubercular drug targets and provides a comprehensive picture of the complexity of *M. tuberculosis* essential cellular functions.

## Introduction

*Mycobacterium tuberculosis* is responsible for approximately 10.4 million new cases of active tuberculosis (TB) infection and 1.4 million deaths annually (1). While TB chemotherapy has a high success rate in curing drug susceptible TB infections, it is challenging, in part because it requires a minimum of 6 months of treatment with drugs associated with adverse reactions. Thus, finding targets for new TB drugs that are more potent than existing drugs is needed (2).

Essential genes, defined as genes indispensable for growth and/or survival, are potential targets for new types of antimicrobial drugs. Gene essentiality can be assessed by targeted gene-disruptions, where genes that cannot be disrupted are typically categorized as being essential. However, such traditional genetic approaches are labor-intensive and not easily adaptable to genome-scale screening. Recent advances in Next-Generation Sequencing (NGS)-based approaches have transformed our ability to examine gene functions in a genome-wide manner. Transposon insertion sequencing (Tn-seq) has been widely used to conduct fitness profiling of gene functions in many bacterial species, including *M. tuberculosis* (3-10). In addition to fitness profiling, a lack of representation of specific transposon insertions within a saturated transposon library has been used to identify essential genes in genome-wide screens (3, 9, 10).

In *M. tuberculosis*, several systematic genome-wide studies, including studies using Tn-seq, have been performed to identify essential genes *in vitro* (7-16) and *in vivo* (4). The gene essentialities determined by Tn-seq studies are accessible through publicly available databases, such as Tuberculist (17), Biocyc (18), and OGEE (19). Most of these gene essentiality data were obtained from Tn-seq studies that were carried out using a defined growth medium that was supplemented with a limited number of nutrients (7-10, 13, 15). Under this growth condition, genes for numerous essential central metabolic pathways can be rendered dispensable through supplementation. Thus, these previous studies have likely miscategorized a large set of genes as essential rather than conditionally essential. For example, pantothenate, an essential precursor in coenzyme A biosynthesis, is not supplemented in most of the commonly used media for *M. tuberculosis*. As a result, *panC*, encoding pantothenate synthetase, the enzyme catalyzing the last step in pantothenate biosynthesis, was defined as an essential gene in these three databases (17-19) despite a previous study showing that the *panCD* double deletion mutant of *M. tuberculosis* H37Rv strain can grow in the presence of supplemental pantothenate (20). Thus, more precise definition of *M. tuberculosis* essential genes on a genome-wide scale, and annotation of genes which are conditionally essential, would improve the usefulness of these databases.

In this study, we developed a defined-nutrient rich medium (Mtb YM rich) that included a variety of nutrient sources for *M. tuberculosis* to ease identification of conditionally essential genes. We used Tn-seq to identify essential genes in Mtb YM rich and minimal medium (Mtb minimal). As expected, essentialities of many genes involved in metabolite biosynthesis pathways were affected by the supplemental nutrients in Mtb YM rich medium. However, we found that essentialities of certain metabolic pathways were unaffected by the absence or presence of relevant nutrient sources. We also found that both growth conditions had some essential genes that were unique to each environment. In addition, we compared essential genes identified by Tn-seq with highly conserved genes that were identified by a comparative genomics analysis and a modified *in silico* metabolic model. These comparisons indicated that essential genes were highly enriched amongst the conserved core genome and that such gene essentiality measurements can be used to refine metabolic models.

## Results and Discussions

### Identification of *M. tuberculosis* essential genes in a chemically defined nutrient-rich media

We developed a defined nutrient-rich medium for *M. tuberculosis* (Mtb YM rich medium) (Fig. 1A, Table S1) by supplementing numerous nutrients into a minimal *M. tuberculosis* growth medium (Mtb minimal medium) (21). We chose supplements based on their known use by various bacterial species including *M. tuberculosis* (7, 12, 20, 22-24). The supplements included several carbon sources, nitrogen sources, co-factors, amino acids, nucleotide bases and other nutrients. Four of the supplements, lipoic acid, nicotinamide, hemin, and ribose, inhibited *M. tuberculosis* H37Rv growth at higher concentrations. Thus, we utilized concentrations of these supplements that did not impair growth. The Mtb YM rich medium supported the *M. tuberculosis* H37Rv growth similar to the commonly used 7H9 medium that contains essential salts and relatively few nutrients (Fig. 1B).

**Figure 1.**
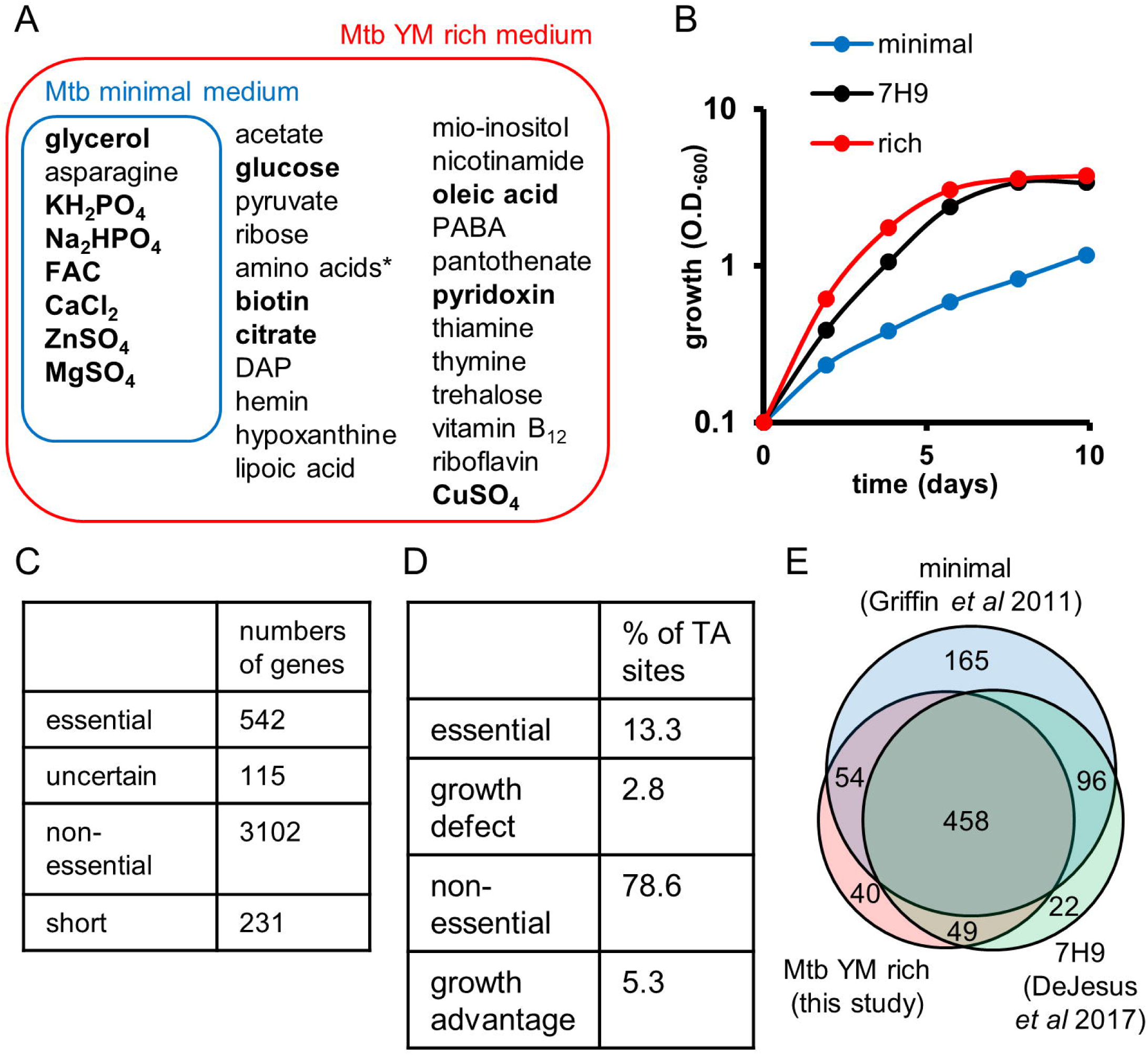
Identification of essential genes in Mtb YM rich medium. (A) Media composition of Mtb minimal and Mtb YM rich. 7H9 media contains glutamic acid and ammonium sulfate in addition to the nutrients shown in bold. DAP, diaminopimelic acid. FAC, ferric ammonium citrate. PABA, *para*-aminobenzoic acid. *, 0.5 % casamino acids and 0.98 mM tryptophan. (B) *M. tuberculosis* H37Rv growth in Mtb YM rich (red), 7H9 (black), and Mtb minimal (blue). (C) Bayesian/Gumbel results for *M. tuberculosis* H37Rv in Mtb YM rich. (D) HMM results for *M. tuberculosis* H37Rv in Mtb YM rich. (E) Venn diagram comparisons of essential genes by Tn-seq in this study and two previous studies (8, 9). This study (red), Griffin *et al* (blue) (8), and DeJesus *et al* (green) (9).

We next generated a library of *M. tuberculosis* H37Rv transposon insertion mutants on Mtb YM rich medium plates. As the *M. tuberculosis* H37Rv genome contains 74,602 TA dinucleotides, we collected at least 150,000 colonies to approach saturation of *himar1* transposon insertion sites in the library. Genomic DNA was isolated from the library. Next, the DNA was sheared, end repaired, sequencing adapters were added, and the transposon adjacent regions were enriched by PCR before massively parallel sequencing. The resultant sequencing data was analyzed using TRANSIT, a recently developed software package for analyzing Tn-seq data (14). TRANSIT contains two statistical methods, Bayesian/Gumbel Method and Hidden Markov Model (HMM), to identify essential genes or essential genomic regions, respectively, in a single growth condition. By using the Bayesian/Gumbel Method, we found that 542 genes were essential for *M. tuberculosis* growth in the Mtb YM rich medium (Fig. 1C, Dataset S1). The Bayesian/Gumbel method assesses gene essentiality based on consecutive sequence of TA sites lacking insertion within a gene. When the analysis did not exceed the significance thresholds, genes were called either short or uncertain by TRANSIT. As a result, 231 genes were called short and 115 genes were called uncertain by this analysis (Fig. 1C). To overcome this issue, we also identified essential genomic regions using an HMM (14). The HMM is based on the read count at a given site and the distribution over the surrounding sites. This analysis identified 13.3 % of TA sites as essential and 2.8 % of TA sites as growth defective (Fig. 1D, Dataset S1). These essential and growth defective regions included 17 short genes (14 essential and 3 growth defective genes) and 21 uncertain genes (10 essential and 11 growth defective genes) whose essentiality could not be assessed by the Bayesian/Gumbel Method. In addition, HMM identified 5 short genes and 16 uncertain genes that contained both essential and non-essential TA sites. These genes are listed in Table S2. In total, we identified 601 genes (542 genes by the Bayesian/Gumbel Method and additional 59 genes by the HMM) as essential genes for *M. tuberculosis* survival in the Mtb YM rich medium. A list of these genes is available through the BioCyc smart table format (https://biocyc.org/group?id=biocyc14-7907-3764257976) (25). These essential genes included known targets for existing anti-tubercular drugs further validating that the Tn-seq assay successfully identified *M. tuberculosis* essential genes (Table 1).

**Table 1.**
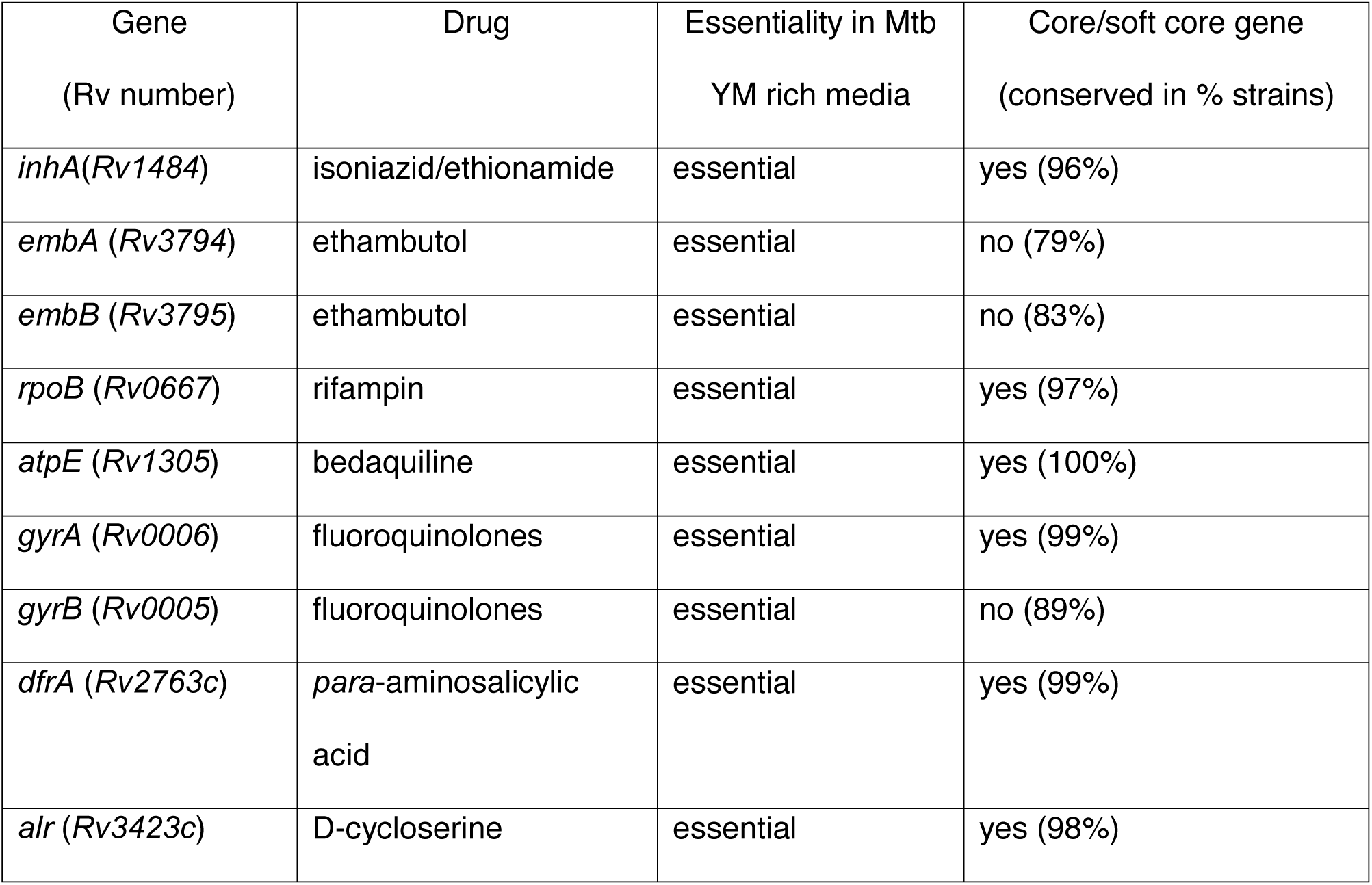
Gene essentialities and *in silico* essentiality predictions of anti-tubercular drug target genes.

| Gene<br>(Rv number) | Drug | Essentiality in Mtb<br>YM rich media | Core/soft core gene<br>(conserved in % strains) |
| --- | --- | --- | --- |
| *inhA* (Rv1484) | isoniazid/ethionamide | essential | yes (96%) |
| *embA* (Rv3794) | ethambutol | essential | no (79%) |
| *embB* (Rv3795) | ethambutol | essential | no (83%) |
| *rpoB* (Rv0667) | rifampin | essential | yes (97%) |
| *atpE* (Rv1305) | bedaquiline | essential | yes (100%) |
| *gyrA* (Rv0006) | fluoroquinolones | essential | yes (99%) |
| *gyrB* (Rv0005) | fluoroquinolones | essential | no (89%) |
| *dfrA* (Rv2763c) | *para*-aminosalicylic<br>acid | essential | yes (99%) |
| *alr* (Rv3423c) | D-cycloserine | essential | yes (98%) |

The list of essential genes identified in this study was also compared with the essential genes identified by the past Tn-seq studies, Griffin et al (8) and DeJesus et al (9) (Fig. 1E and Table S3). In general, our result was largely consistent with the past studies and a total of 458 genes were identified as essential in all three studies (Fig. 1E) (https://biocyc.org/group?id=biocyc14-7907-3764424447). Since Mtb YM rich medium used in this study was supplemented with numerous nutrients, many genes that were related to biosynthetic pathways of the supplemented nutrients (e.g. amino acids, pantothenate, purine, flavin and others) were not essential in our study while the past studies categorized these genes as essential. We also identified genes that were only essential in our study but not in other studies. We identified some of these genes as conditionally essential in Mtb YM rich media (described below, Fig. 2, and Table S4). Of note, our study could not detect several genes that were highly expected to be essential and identified as essential in DeJesus study. These genes included short genes such as several ribosomal genes and *folK*. This likely was because DeJesus study used a more saturated transposon library (fourteen replicates vs two replicates).

**Figure 2.**
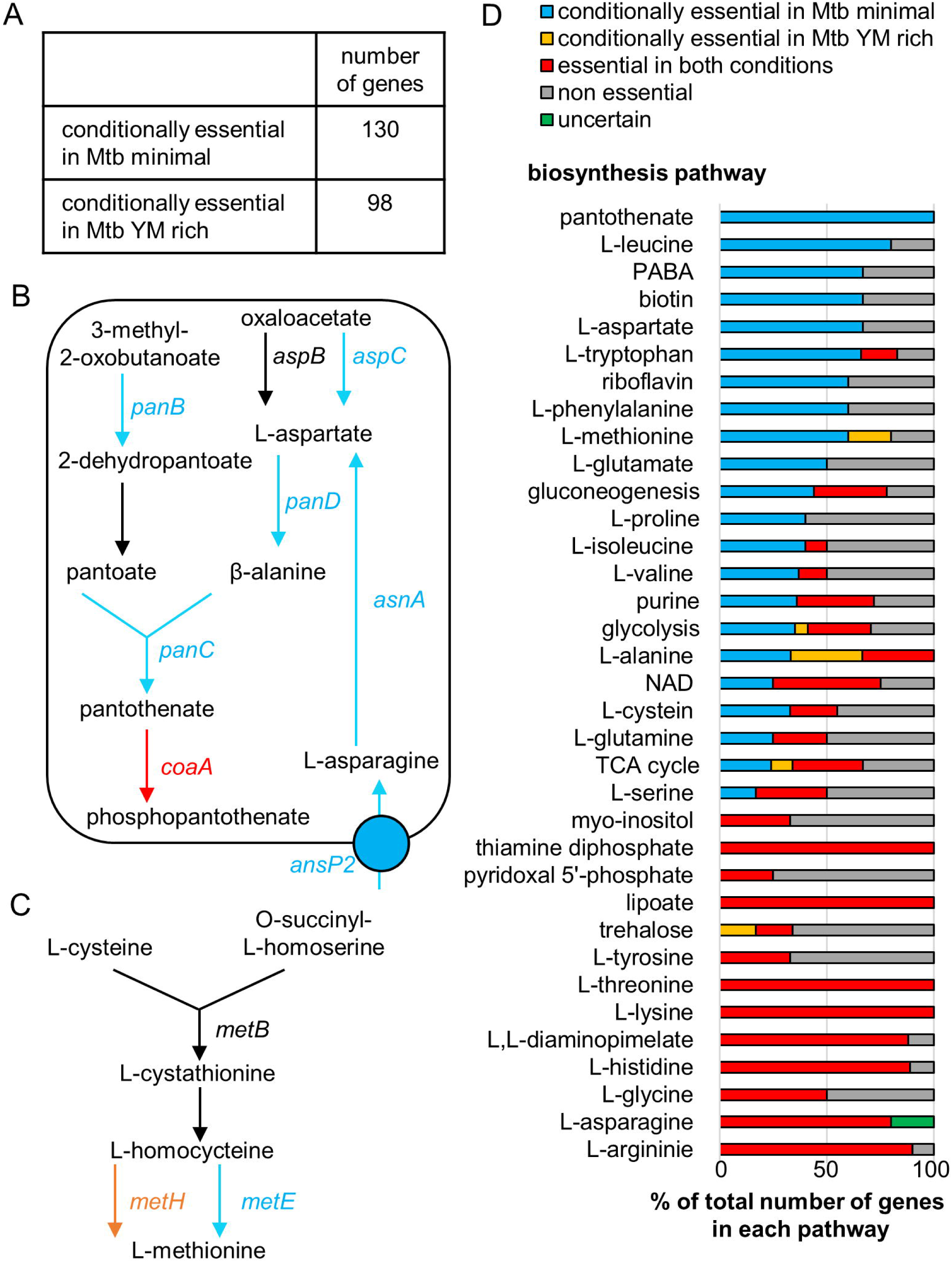
Identification of conditionally essential genes in Mtb YM rich and Mtb minimal media. (A) Results for comparative analysis between Mtb YM rich and Mtb minimal media. Numbers of conditionally essential genes (adjusted p-value <0.05) were identified by the resampling method in TRANSIT (14). (B, C) Essentialities of genes in pantothenate biosynthesis pathway (B) and methionine biosynthesis pathway (C). Genes in red color (essential in both Mtb YM rich and Mtb minimal), genes in blue color (essential in Mtb minimal but not essential in Mtb YM rich), genes in orange color (essential in Mtb YM rich but not essential in Mtb minimal) genes in black color (non-essential). Information on genes in each pathway was obtained from BioCyc database (18). (D) Ratio of essential genes in each metabolic pathway. All genes are listed in Table S5. NAD, Nicotinamide adenine dinucleotide. PABA, *para*-aminobenzoic acid. purine, 5-aminoimidazole ribonucleotide.

### Comparison of *M. tuberculosis* essential genes between Mtb YM rich medium and minimal nutrient medium

Because gene essentiality can be affected by the external environment (26), in particular by nutrient availability, we next compared gene essentiality between Mtb YM rich medium and Mtb minimal medium. Similar to the transposon library generated on Mtb YM rich medium plates, we generated *M. tuberculosis* H37Rv transposon libraries on Mtb minimal medium plates and collected at least 150,000 colonies to create a saturated library. The resultant sequencing data was compared with the Mtb YM rich plate data by using a permutation test-based method to identify genes with statistically significant differences in transposon insertion count (Fig. 2A, Table S4). We found that 130 genes were conditionally essential in the Mtb minimal medium compared to the Mtb YM rich medium (https://biocyc.org/group?id=biocyc13-7907-3706551227).

As anticipated, many of the conditionally essential genes that were identified corresponded to the differences in nutrient composition between Mtb minimal and Mtb YM rich. For example, Mtb YM rich medium contained L-aspartate and pantothenate. Thus, the genes in L-aspartate and pantothenate biosynthesis pathways were dispensable in Mtb YM rich medium (Fig. 2B). However, Mtb YM rich medium did not contain metabolites located downstream of pantothenate. Consequently, these downstream genes (e.g. *coaA*, encoding pantothenate kinase) were essential in both Mtb YM rich and Mtb minimal media (Fig. 2B). This observation is consistent with a previous report that the *M. tuberculosis panCD* double deletion mutant strain can grow in the presence of pantothenate (20), and also suggested that single gene disruptions in the pantothenate biosynthesis pathway (*panB, panC*, and *panD*) show a similar pantothenate auxotrophic phenotype as the *panCD* double deletion mutant strain. Another example was the asparagine transporter gene *ansP2* (27) that was conditionally essential in Mtb minimal medium (Fig. 2B) presumably because Mtb minimal medium contained arginine as the sole nitrogen source whereas Mtb YM rich medium contained multiple nitrogen sources (Fig. 1A).

Unexpectedly, we also identified 98 genes that were conditionally essential in Mtb YM rich media compared to the Mtb minimal medium (https://biocyc.org/group?id=biocyc13-7907-3710604059). Such genes included *ponA1* and *ponA2*, encoding penicillin binding proteins (PBPs) involved in cell wall peptidoglycan (PG) biogenesis. It was previously shown that *ponA1* and *ponA2* are only essential *in vivo* but not during growth in culture medium (28, 29). Thus, one of nutrients that is uniquely present in Mtb YM rich medium might be also present *in vivo* and may be responsible for the *in vivo* fitness defect of the *ponA1* mutant. LdtB is one of the major L,D-transpeptidases (Ldts) that is also involved in PG biogenesis. *ldtB* was also conditionally essential in Mtb YM rich medium. Interestingly, several genes that were conditionally essential in the absence of *ponA1, ponA2*, or *ldtB* (e.g. *Rv1086, Rv1248c, Rv3490, treS, otsA*) were also conditionally essential in Mtb YM rich medium suggesting Mtb YM rich medium negatively affects PG biogenesis in *M. tuberculosis* (30). We also found the *metH*, encoding one of the two methionine synthases, was only essential in Mtb YM rich medium whereas *metE*, encoding the other methionine synthase, was only essential in Mtb minimal medium (Fig.2C). The observed conditional essentialities were consistent with a previous study that *M. tuberculosis metE* expression was inhibited in the presence of vitamin B_12_ by a *metE* B_12_ riboswitch, and the vitamin B_12_-dependent MetH is predominantly used when vitamin B_12_ is available (31). These findings confirmed that the conditionally essential genes identified by Tn-seq in this study are consistent with previous findings (25).

The supplementation of nutrients in the Mtb YM rich medium could potentially subvert the need for enzymes in at least 35 metabolic pathways (Fig. 1A, Fig. 2D, and Table S5). We found fewer gene essentialities in 22 of these pathways in Mtb YM rich medium, suggesting that *M. tuberculosis* can functionally utilize these nutrients. Notably, many genes in these pathways were identified as essential genes by the previous studies (8, 13) and listed as essential genes in public database such as Tuberculist (17) and BioCyc (18). Thus, our results revealed that these genes are essential only in the absence of the corresponding nutrients. In contrast, we also found that essentiality of genes in 13 other metabolic pathways were not altered by nutrient supplementation (Fig. 2D). Previous studies have demonstrated that some auxotrophic mutants, such as those for L-arginine, L-lysine, and inositol, require supplementation with the corresponding nutrient at a relatively high concentration to support their growth (32-34). Certain auxotrophic mutants are also known to show growth or survival defects even in the presence of the corresponding nutrient (35, 36). These results were consistent with previous findings that essentiality of some central metabolic pathways could not be bypassed in pathogenic mycobacteria (12) and also provide a more comprehensive understanding of the nutrient utilization capacity of *M. tuberculosis*.

### Identification of highly conserved genes in the *M. tuberculosis* complex

Essential genes that were identified by Tn-seq were further interrogated through comparative genomic analysis in order to see whether essentiality correlated with high levels of sequence conservation within the *M. tuberculosis* complex. A total of 226 complete genome sequences of *M. tuberculosis* complex were available from the PATRIC database (37). We excluded non-pathogenic strains (e.g. *M. tuberculosis* H37Ra and *Mycobacterium bovis* BCG) and used 199 genome sequences for comparative genomic analysis (Fig. 3A, Table S6). We identified total of 17,813 genes from 199 strains of the *M. tuberculosis* complex (Fig. 3B and C). Among these genes, 2, 206 genes (1,030 core genes and 1,176 soft core genes) were highly conserved in most of the strains (≥ 95 % of strains).

**Figure 3.**
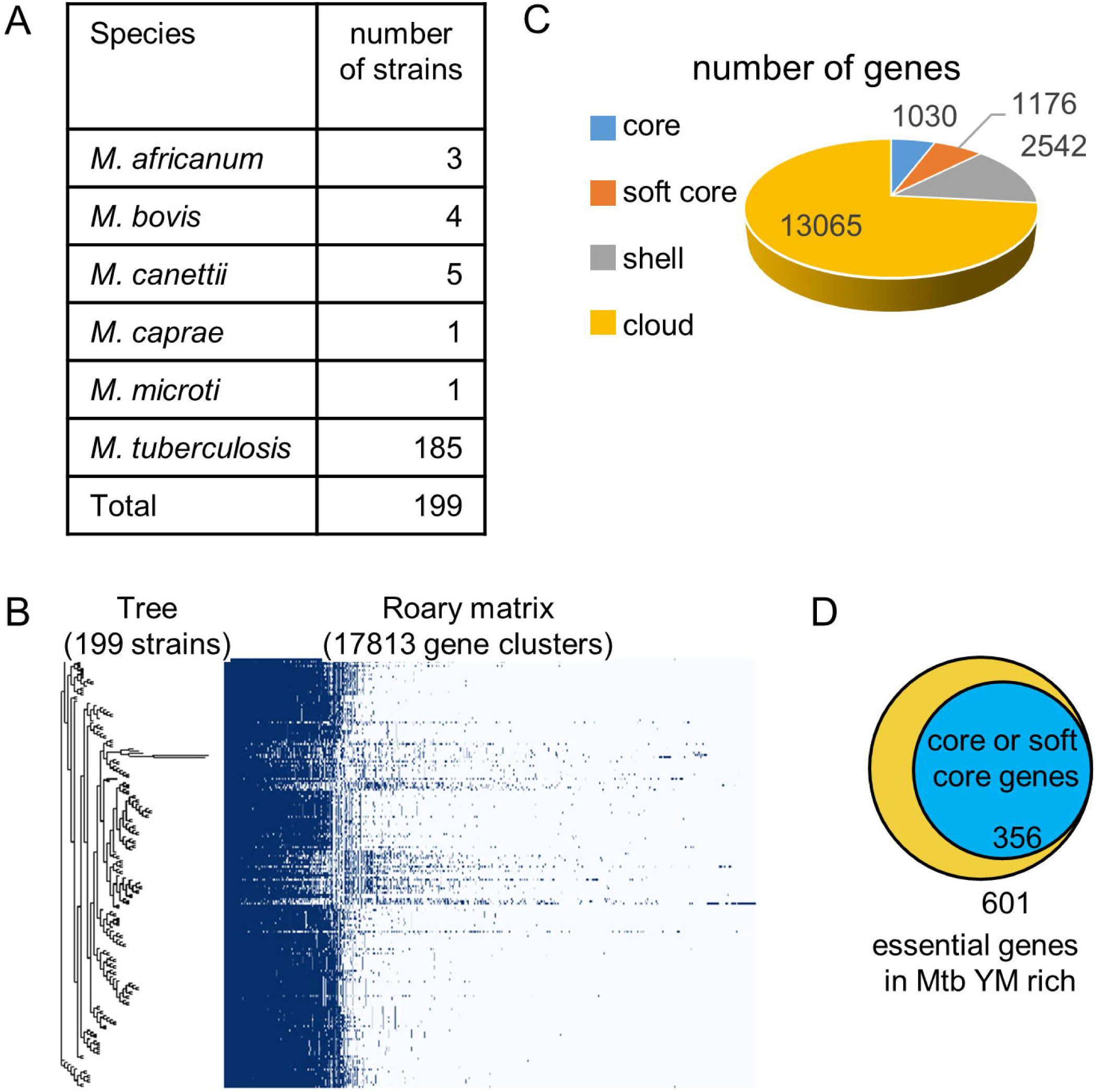
Identification of highly conserved essential genes among virulent *M. tuberculosis* complex. (A) Strains used for comparative genomic analysis. (B) Results for pan-genome analysis by Roary. (C) Number of highly conserved genes among virulent *M. tuberculosis* complex. Core genes; conserved in >99 % strains, soft core genes; conserved in 95-99 % strains, shell genes; conserved in 15-95% strains, cloud genes; conserved in 0-15 % strains. (D) Number of genes that are essential for *M. tuberculosis* H37Rv survival in Mtb YM rich and highly conserved among virulent *M. tuberculosis* complex strains.

Our Tn-seq analysis identified 601 genes as essential in Mtb YM rich medium (Dataset S1, Table S2 and smart table). Among them, we confirmed at least 60 % of the essential genes (356 genes) were included in the list of core/softcore genes (Fig. 3D) (https://biocyc.org/group?id=biocyc14-7907-3764449513). Of note, not all of the target genes for existing anti-tubercular drugs were categorized as core/softcore genes (Table 1).

### Prediction of *M. tuberculosis* essential genes *in silico*

Genome-scale metabolic models have been used to computationally simulate a range of cellular functions (38). We utilized a genome-scale model of *M. tuberculosis*, iSM810, to predict gene essentiality *in silico* using flux balance analysis (FBA) (39). The principles for FBA have been described previously (40). Briefly, a metabolic network is represented by a stoichiometric matrix that contains reactions and metabolites. A biomass reaction is defined based on experimentally determined amounts of specific metabolites required for cellular growth. The stoichiometric matrix can be converted into a system of linear equations. FBA then solves this system by optimization, typically of the biomass reaction. Constraints on reaction fluxes can be used to simulate metabolite availability or enzyme presence. For example, a reaction’s uptake flux is constrained to zero when the simulated environment does not contain the associated metabolite. Reaction fluxes that simulate metabolite uptake within iSM810 were used to define *in silico* growth media. Uptake was not bounded for metabolites present in the selected growth medium, however; not all media components were represented within the iSM810 transport reactions (Table S7). Genes were assessed for essentiality by systematically closing the flux on each reaction within iSM810 and assessing simulated biomass production on each medium using FBA. Genes were defined essential if biomass production after knockout was <1e-10.

iSM810 includes 810 metabolic genes (including 1 orphan gene) and 938 metabolic reactions (39). Among the 810 genes in iSM810, our FBA analysis predicted that 159 genes were essential in Mtb YM rich medium and 221 genes were essential in Mtb minimal medium (Fig. 4A, Table S7). We then compared gene essentiality prediction by FBA with essential genes identified by Tn-seq (Fig. 4B). We found that the sensitivity of the FBA based gene essentiality prediction was low as there were a number of genes that were predicted to be essential by FBA but not identified as essential genes by Tn-seq (Fig. 4B) (https://biocyc.org/group?id=biocyc14-7907-3764439908). For instance, we found that genes related to riboflavin biosynthesis were non-essential in Mtb YM rich medium by Tn-seq analysis, but were essential *in silico*. We examined why iSM810 could not accurately predict the essentiality of genes in the riboflavin biosynthesis pathway and found that the model lacked a transport reaction for riboflavin (Table S7). Similarly, we found that the model lacked transport reactions for vitamin B_12_, *para*-aminobenzoic acid (PABA), H_2_O, and myo-inositol. To investigate whether the addition of these transport reactions could improve gene essentiality prediction by iSM810, we added these transport reactions to the model. These changes fixed multiple mis-matches between iSM810 and Tn-seq, by causing the model to determine that genes related to thiamine biosynthesis and riboflavin biosynthesis were non-essential in Mtb YM rich medium (Fig. 4A and B, Table S7). Allowing PABA uptake caused no changes. After adding transport reactions, the growth rate prediction in Mtb YM rich medium increased from 0.055 g/L/day to 0.0876 g/L/day.

**Figure 4.**
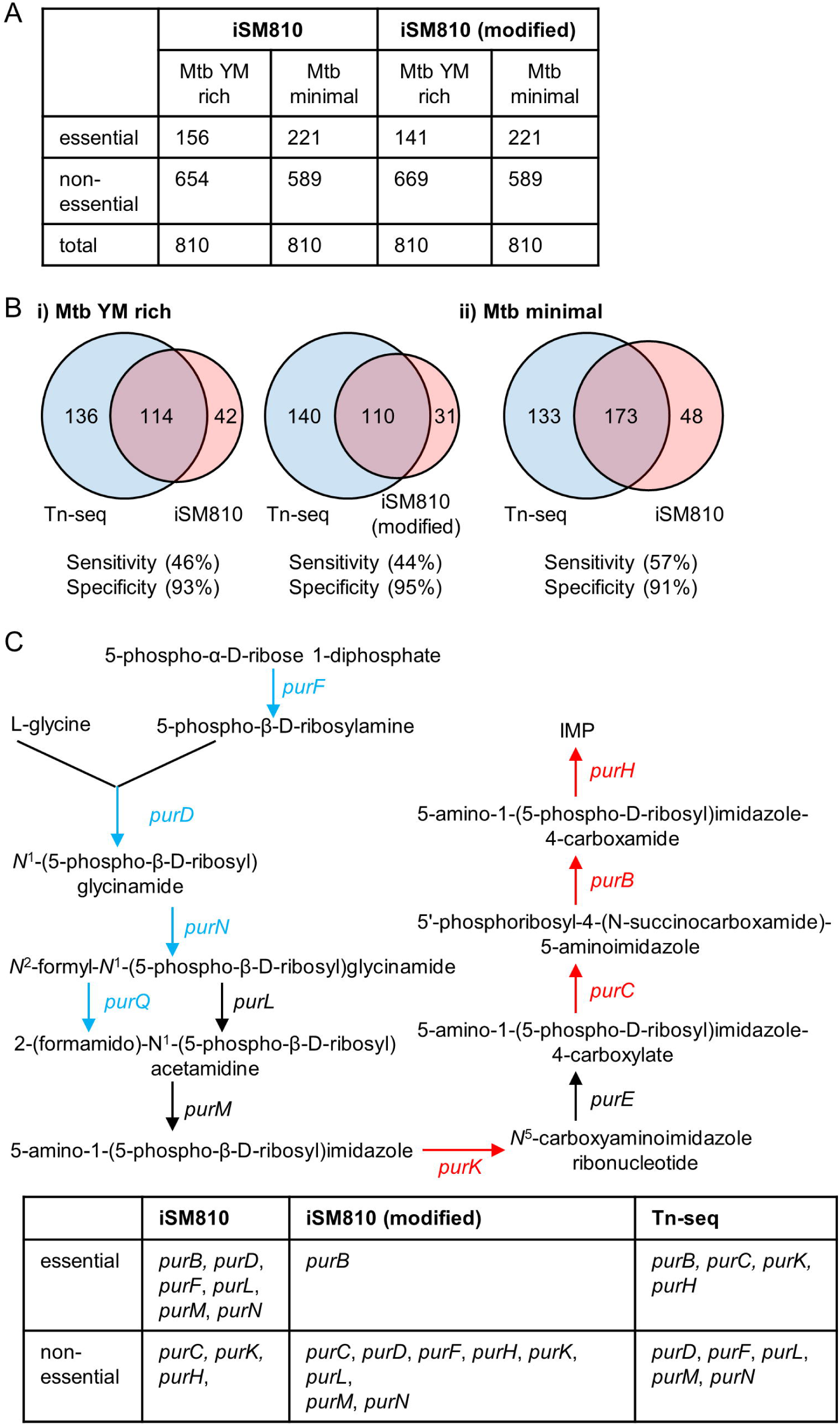
Comparison of Tn-seq identified essential genes and *in silico* predicted essential genes. (A) Results for *in silico* gene essentiality prediction by iSM810 and iSM810 (modified). (B) Comparison of Tn-seq identified essential genes (blue) and iSM810 predicted essential genes (red). (C) Comparison of Tn-seq identified essential genes and iSM810 predicted essential genes in purine biosynthesis pathway. Genes in red color (essential in both Mtb YM rich and Mtb minimal), genes in blue color (essential in Mtb minimal but not essential in Mtb YM rich), genes in black color (non-essential). Information on genes in each pathway was obtained from BioCyc database (18).

Unlike the Tn-seq results, the FBA predicted that all genes essential in Mtb YM rich medium were also essential in Mtb minimal and failed to predict any genes such as *metH* that were only essential in Mtb YM rich (Table S7). This is expected given the lack of gene regulation implicit in our FBA modeling.

Comparing the sets of essential genes experimentally identified by Tn-seq to those computationally predicted *in silico* highlighted the limitations of current *M. tuberculosis* metabolic models. For example, we noticed that genes located downstream of purine biosynthesis pathway were identified as essential by Tn-seq in both Mtb YM rich and Mtb minimal media (Fig. 4C). However, there were mismatches between iSM810 and Tn-seq. Among these genes, *purC* is the only gene that was characterized by targeted gene deletion analysis in *M. tuberculosis* (41). The *purC* gene was difficult to delete (41) and the *purC* deletion mutant strain of *M. tuberculosis* showed a notable growth defect compared to the parent strain even in the presence of hypoxanthine (22). This observation could explain why the *purC* gene was identified as essential by Tn-seq in Mtb YM rich. However, the other genes in the purine biosynthesis pathway have not been characterized by targeted gene deletion studies. Thus, future studies are necessary to investigate essentiality of *purK, purB*, and *purH* by targeted gene deletion.

We also identified a potential avenue for improvement of the genome-scale model of *M. tuberculosis* by comparing Tn-seq identified essential genes and to those predicted *in silico.* We found that many genes in known essential metabolic pathways were not predicted to be essential (e.g. glycolysis, folate metabolism, mycolate biosynthesis, ATP biosynthesis, and several amino acid biosynthesis pathways) (https://biocyc.org/group?id=biocyc14-7907-3764451955). Examination of the biomass function in iSM810 revealed that several known essential metabolites are not connected to the biomass function in iSM810. For example, folate is not connected to the biomass function and this explains why addition of a PABA transport reaction did not affect any gene essentiality predictions. None of ATP synthase genes in iSM810 (*atpA, atpB, atpD, atpE, atpG*, and *atpH*) was predicted as essential because these were linked to the same reaction using “or” Boolean logic, meaning the presence of one gene was sufficient to allow the reaction to proceed. Thus, further improvement of iSM810 for more accurate prediction of essential genes could be achieved by connecting such essential metabolites to the biomass function.

### Conclusions

In this study, we utilized functional genomics and comparative genomics approaches to identify *M. tuberculosis* essential genes. We identified distinct sets of essential and conditionally essential genes from different approaches and different growth conditions. In addition, comparison of essential genes identified by functional genomics approach to *in silico* predicted essential genes highlighted current gaps in knowledge regarding *M. tuberculosis* metabolism. Our study provides a promising platform to shed new light on essential cellular functions in *M. tuberculosis* that can lead to the discovery of novel targets for antitubercular drugs.

## Materials and methods

### Media and growth conditions

*M. tuberculosis* H37Rv was grown aerobically at 37 °C in Middlebrook 7H9 medium supplemented with oleate-albumin-dextrose-catalase (OADC; 10%, vol/vol), glycerol (0.2%, vol/vol), and tyloxapol (0.05%, vol/vol) unless otherwise noted. Mtb minimal medium (0.2%, vol/vol glycerol as a sole carbon source) (21) and Mtb YM rich medium (Table S1) agar plates were used to generate *M. tuberculosis* H37Rv transposon libraries. Tyloxapol (0.05%, vol/vol) were added to the both Mtb minimal and Mtb YM rich agar plates.

### Construction of saturated transposon libraries of *M. tuberculosis*

Transposon mutagenesis were performed as previously described (42). Mycobacteriophage phAE180 (42) was used to transduce a mariner derivative transposon Tn5371 (43) to *M. tuberculosis* H37Rv that was grown until mid-log growth phase (OD_600_ 0.5). Transduced *M. tuberculosis* H37Rv were spread on Mtb minimal plate and Mtb YM rich plate and incubated at 37 °C for 2-3 weeks. To generate a transposon library in saturated size, at least 150,000 colonies were collected from each plate. Each transposon library was aliquoted and stored at −80 °C. Each transposon library was generated in duplicates.

### Tn-seq

Genomic DNA (gDNA) was prepared from each sample as previously described (44). gDNA was then fragmented using an S220 Acoustic DNA Shearing Device (Covaris). After shearing, adapters were added using an Illumina TruSeq Nano DNA library prep kit according to the manufacturer’s instructions. Transposon junctions were amplified by using a transposon specific primer Mariner_1R_TnSeq_noMm (TCGTCGGCAGCGTCAGATGTGTATAAGAGACAG<u>CCGGGGACTTATCAGCCAACC</u>) and p7 primers (CAAGCAGAAGACGGCATACGAG) with HotStarTaq Master Mix Kit (Qiagen) with the following PCR condition (94 °C for 3 min, 30 cycles of 94 °C for 30 seconds, 65 °C for 30 seconds, and 72 °C for 60 seconds, 72 °C for 10 minutes). The transposon junction enriched sample was diluted 1:50 with water and then amplified to add the flow cell adapter and i5 index to the enriched Tn-containing fragments using the following primers: i5 indexing primer [XXXXXXX denotes the position of the 8 bp index sequence): AATGATACGGCGACCACCGAGATCTACACXXXXXXXXTCGTCGGCAGCGTC p7 primer: CAAGCAGAAGACGGCATACGAG

The amplification reaction was as follows: 5 µl template DNA (from PCR 1), 1 µl nuclease-free water, 2 µl 5x KAPA HiFi buffer (Kapa Biosystems), 0.3 µl 10 mM dNTPs (Kapa Biosystems), 0.5 µl DMSO (Fisher Scientific), 0.2 µl KAPA HiFi Polymerase (Kapa Biosystems, 0.5 µl i5 indexing primer (10 µM), 0.5 µl p7 primer (10 µM). Cycling conditions were: 95°C for 5 minutes, followed by 10 cycles of 98°C for 20 seconds, 63°C for 15 seconds, 72°C for 1 minute, followed by a final extension at 72°C for 10 minutes. Amplification products were purified with AMPure XP beads (Beckman Coulter) and the uniquely indexed libraries were quantified using a Quant-IT PicoGreen dsDNA assay (Thermo Fisher Scientific) and the resulting fragment size distribution was assessed using a Bioanalyzer (Agilent Technologies). The resultant Tn-seq library was sequenced using a HiSeq 2500 HO, 125 bp PE run using v4 chemistry (Illumina).

### Tn-seq analysis

Sequence reads were trimmed using CutAdapt (45). We first trimmed sequence reads for transposon sequences (CCGGGGACTTATCAGCCAACCTGT) at the 5’ ends. Reads that did not contain transposon sequence at the 5’ end were discarded. After the 5’ trimming process, all the sequence reads begin with TA. We then trimmed sequence reads for adaptor sequences ligated to 3’end (GATCCCACTAGTGTCGACACCAGTCTC). After the trimming, we discarded the sequence reads that were shorter than 18 bp. The default error rate of 0.1 was used for all for all trimming processes.

The trimmed sequence reads were mapped (allowing 1 bp mismatch) to the *M. tuberculosis* H37Rv genome (GenBank: AL123456.3) using Bowtie 2 (46). The number of reads at each TA site were counted and converted to the .wig format, the input file format for TRANSIT (14) using a custom Python script (S1 Appendix). Subsequent statistical analysis for gene essentiality (Bayesian/Gumbel Method, HMM, and resampling method) were performed using TRANSIT (version, 2.0.2) (14).

The Bayesian/Gumbel Method determines posterior probability of essentiality of each gene (shown in zbar column in Dataset S1). When the value is 1 or near 1 within the threshold, gene is called essential. When the value is zero or near zero the threshold, gene is called non-essential. When the value is between the two threshold, neither near zero nor 1), gene is called uncertain. When the value is −1, gene is called small because it is considered that gene is too small to determine posterior probability of essentiality. Thus, we analyzed essentialities of small and uncertain genes by HMM. All essential genes identified from uncertain or small genes were listed in Table S2.

Total 601 genes (https://biocyc.org/group?id=biocyc14-7907-3764257976) essential for *M. tuberculosis* survival in the Mtb YM rich medium (542 genes by the Bayesian/Gumbel Method and additional 59 genes by the HMM) were used for the comparisons to two past Tn-seq studies (8, 9). List of essential genes identified by past studies were obtained from Dataset S1 in ref (8) and ref (9).

### Flux Balance Analysis (FBA)

FBA solutions were obtained using a simulated environment designed to mimic the Mtb YM rich medium designed for this study. This was done by altering uptake boundaries to match the concentration of each metabolite. Most metabolites present in the media were given unlimited boundaries, because these metabolites were not expected to be limiting, and also because they were present at an undefined concentration in the Mtb YM rich medium, due to their source being casamino acids. Metabolites added to the Mtb YM rich medium in known concentrations were bounded in FBA at those concentrations.

The iSM810 model contains 938 metabolic reactions and 810 genes (including 1 orphan gene) (39). The biomass reaction originally described for iSM810 was chosen to define growth. FBA was performed using the COBRA Toolbox Matlab package (47, 48). The unconstrained uptake fluxes were set to ‘1’. Gene essentiality was assessed using the COBRA Toolbox single gene deletion function in Matlab. Through single gene deletion, reactions associated with each gene were systematically closed and the model was optimized for biomass production. Any biomass accumulation > 1e-10 (which could occur due to numerical errors) was defined as growth and the gene was classified as non-essential. Biomass accumulation <1e-10 resulted in a gene being called essential. All FBA optimizations were done using the Gurobi Optimizer 7.0 software under the free academic license (49).

### Comparative genomic analysis

Following comparative genomic analysis was carried out as previously reported (50). In brief, the pan and core genomes were defined using Roary software (51). Complete and draft genome sequences of pathogenic strains (non-highlighted strains were used and obtained from PATRIC database (1st October, 2017) summarized in Table S6 were reannotated to generate gff3 files using PROKKA ver. 1.1.12 software (52) and to include annotation of a reference strain H37Rv. Homologous proteins (i.e. protein families) were clustered using the CD-Hit and MCL algorithms. The BLASTp cut-off value was set at 95%. The number of core- and pan-genome protein families were estimated via genome sampling up to the number of input genomes at the default setting in Roary (Dataset S2).

## Acknowledgments

We thank Nicholas D. Peterson and Dr. Igor Libourel for their assistance in initial conception and design of the study. This study was supported by funds from the Minnesota Partnership for Biotechnology and Medical Genomics (ML2012, chapter 5, article 1, section 5, subdivision 5e to A.D.B), the American Lung Association (to A.D.B), the National Institutes of Health (GM121498, to W.R.H. and AI123146 to A.D.B.) the Japan Agency for Medical Research (AMED; project number 17fk0108116h0401to FM), Kakenhi (18K19674 and 16H05501 to F.M).

## Supporting Information Legends

**Table S1. Composition of Mtb YM rich, Mtb minimal media.**

**Table S2 List of short and uncertain genes that were identified as essential by TRANSIT HMM.**

**Table S3 List of essential genes identified in this study and past studies.**

**Table S4 Comparison of essential genes between Mtb YM rich and Mtb minimal media identified by Tn-seq and analyzed by TRANSIT re-sampling method.**

**Table S5 Comparison of essential metabolic pathways between Mtb YM rich and Mtb minimal media.**

**Table S6 Complete and draft genome sequences of virulent *M. tuberculosis* complex used for comparative genomic analysis.**

**Table S7 *in silico* growth predictions of a single gene knockout mutant strains of *M. tuberculosis*.**

**Dataset S1 Analysis of Tn-seq gene essentiality analysis by TRANSIT Bayesian/Gumbel Method and Hidden Markov Model.**

**Data S2. Presence or absence of genes within complete genomes of the *Mycobacterium tuberculosis* complex analyzed by Roary.**

**Appendix S1. Custom Python script used in this analysis.**

## Notes

#### Summary of Updates

Main text, all figures, supplemental files updated.

